# Rapid adaptation and increased genetic parallelism in experimental metapopulations of *Pseudomonas aeruginosa*

**DOI:** 10.1101/2024.10.28.620660

**Authors:** Partha Pratim Chakraborty, Rees Kassen

**Affiliations:** University of Ottawa, Ottawa, Ontario, Canada; Mcgill University, Montreal, Quebec, Canada

## Abstract

Natural populations are often spatially structured, meaning they are best described as metapopulations composed of subpopulations connected by migration. We know little about how the topology of connections in metapopulations impacts adaptive evolution. Topologies that concentrate dispersing individuals through a central hub can accelerate adaptation above that of a well-mixed system in some models, however empirical support is lacking. We provide evidence to support this claim and show acceleration is accompanied by high rates of parallel evolution resulting from a reduced probability that rare beneficial mutations are stochastically lost. Our results suggest metapopulation topology can be a potent force driving evolutionary dynamics and patterns of genomic repeatability in structured landscapes such as those involving the spread of pathogens or invasive species.

## Main text

Natural populations are often spatially structured, meaning they are geographically subdivided and connected by migration. Infectious diseases transmit through contact networks among hosts(*1*), species ranges can often be discontiguous, with peripheral populations being connected via dispersal to a more well-connected central population(*2, 3*), and patients (along with their colonizing pathogens) regularly moving among wards in a hospital(*4*). How metapopulation topology, the arrangement of connections among subpopulations, influences the dynamics of adaptation is not well understood. The central issues concern the impact of topology on rates of adaptation, the time to generate and fix beneficial mutations(*5–7*), and the population genetic mechanisms responsible.

The conventional view, based on models of migration-selection balance in populations with non-overlapping generations where reproduction involves offspring replacing parents *en masse*, is that the topology of connections among subpopulations has little influence on rates of adaptation(*8–11*). Migration in these models tends to slow but not prevent adaptation across a metapopulation relative to a well-mixed system when the rate of selection, *s*, exceeds that of migration, *m*. Rates of adaptation in spatially structured populations can be slowed further by clonal interference, the competition for fixation among independently arising beneficial mutations(*12*). This can be particularly true in large populations because mutation supply rates, being the product of population size and mutation rate, are often high in large populations.

By contrast, topology can strongly modulate the dynamics of adaptation in finite populations where individuals reproduce in overlapping generations, either decreasing or in some cases increasing rates of adaptation relative to a well-mixed system(*13–16*). Much attention has been paid to how ‘star’ topologies, which are comprised of a central ‘hub’ connected to peripheral ‘leaf’ subpopulations, accelerate adaptation because this result is unexpected from standard models and may reasonably capture features of many real-world scenarios such as the edge of species ranges or patient transfers from regional care centers to and from large urban hospitals(*17, 18*). Theory predicts that accelerated rates of adaptation in a star relative to a well-mixed population occurs when mutation supply rates (the product of population size, *N*, and mutation rate, *m*) are << 1 (*5, 6*) because rare beneficial mutations become concentrated in the hub and so are less likely to be stochastically lost when rare, a phenomenon called drift loss.

Empirical evidence on the impact of topology on rates of adaptation is limited. In microbial evolution experiments, where population sizes are usually very large, various forms of spatial structure almost invariably slow down adaptation relative to a well-mixed condition (*19–24*), consistent with predictions from standard theory. The one exception is a study where faster adaptation occurred in structured compared to unstructured populations, a result attributed to the ability of the structured population to explore higher fitness peaks in a rugged fitness landscape (*25*). The one study to evaluate the impact of topology *per se* on adaptation across a range of population sizes tracked an initially rare beneficial mutation introduced into one peripheral ‘leaf’ through both ‘star’ and well-mixed metapopulations(*26*). The mutation spread more rapidly through star metapopulations than well-mixed populations when migration rates were low (*m* < 0.01%) and this difference could be exaggerated by reducing population sizes (from *N ∼* 10^7^ to 10^5^ cfu/ml) and biasing migration from leaves toward the hub. The mechanism responsible for the accelerated rate of spread was the avoidance of drift loss: beneficial mutations rising to high frequency in their introduced subpopulation become concentrated in the hub via migration, allowing them to spread rapidly to other leaves(*26*).

Is acceleration possible under more prolonged and open-ended evolution, where mutations arise naturally at any location across the metapopulation and compete with each other for fixation through clonal interference? Evolutionary graph theory, which is based on reproduction of individuals in overlapping generations, suggests the answer depends on the relative rates of drift loss and clonal interference associated with migration; only when clonal interference is sufficiently low to allow rare beneficial mutations to gain access to the hub do we expect to see acceleration(*5, 6, 27*). To test this prediction, we tracked rates of adaptation of the opportunistic pathogen *Pseudomonas aeruginosa* in the presence of subinhibitory concentrations of the fluoroquinolone antibiotic ciprofloxacin for approximately 100 generations in 4-patch star and well-mixed populations across a range of mutation supply rates. We manipulate mutation supply rates by adjusting both the effective population size and migration rate (Fig. 1; see materials and methods). Each treatment combination is replicated 8 times.

**Figure 1:**
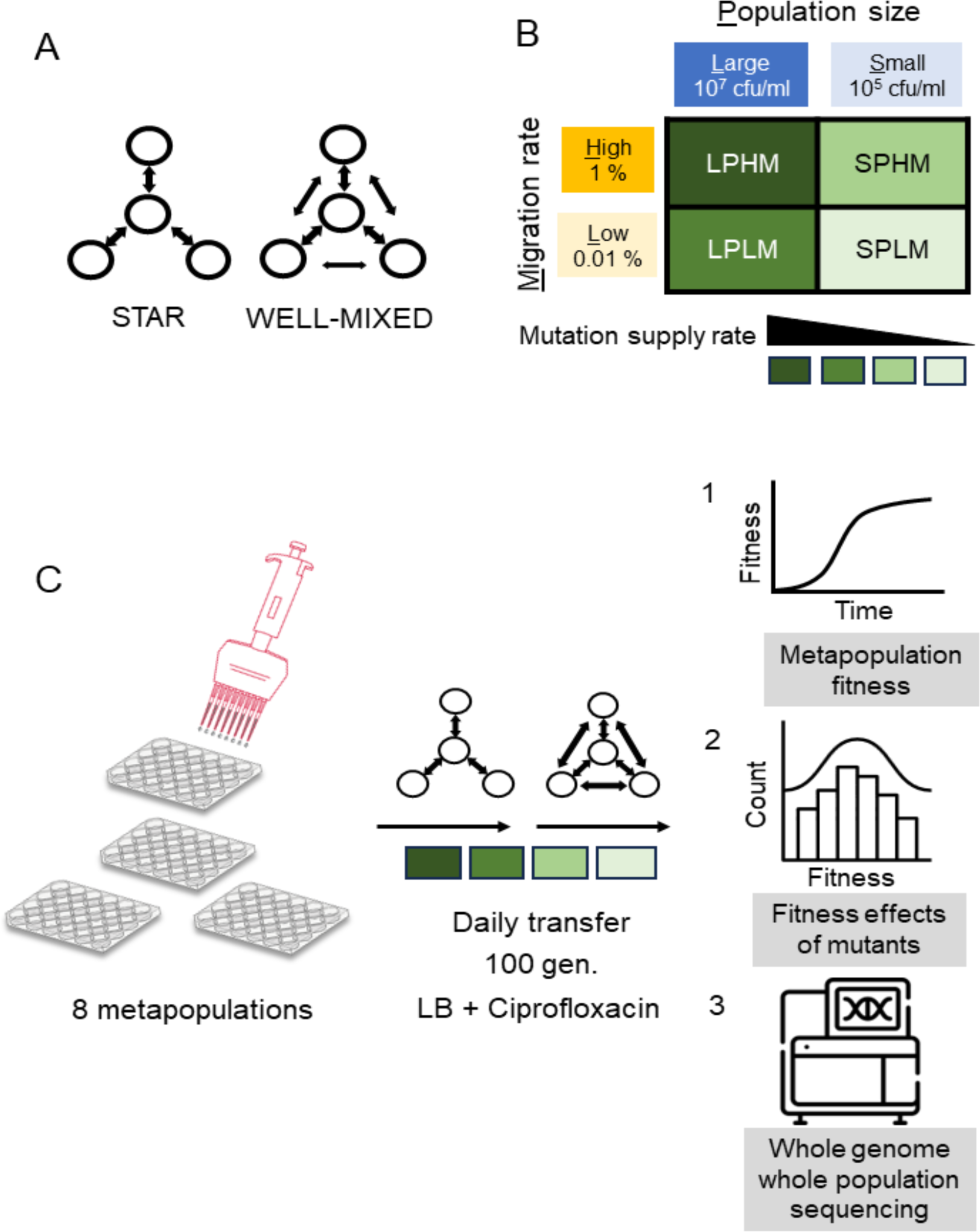
Design for *de novo* evolution experiment. (a) Two topologies, star or well-mixed, constructed among four subpopulations. Arrows depict dispersal routes among subpopulations (circles). (b) Four combinations of mutation supply rates achieved by manipulating both effective population sizes of the subpopulations and the migration rates among the subpopulations. (c) Experimental evolution setup and subsequent assays performed.

Our results are shown in figures 2 and 3. Fitness, measured as both growth rate (*r*) and stationary phase density (*K*) relative to the ancestral strain, increased faster in star metapopulations relative to well-mixed populations when effective population sizes were small (main effect of network, p=0.06 and p<0.01 for relative *r* and *K*, respectively), irrespective migration rate (relative *r*: p=0.73 and relative *K*: p = 0.97). Notably, there was no effect of topology on rates of fitness increase at large effective population sizes (main effect of network treatment for relative *r*: p = 0.95 and relative *K*: p = 0.60) across the two migration rates (relative *r*: p = 0.42 and relative *K*: p = 0.51). These results lend support to the predictions from evolutionary graph theory that star topologies can accelerate adaptation when mutation supply is low but not high.

**Figure 2:**
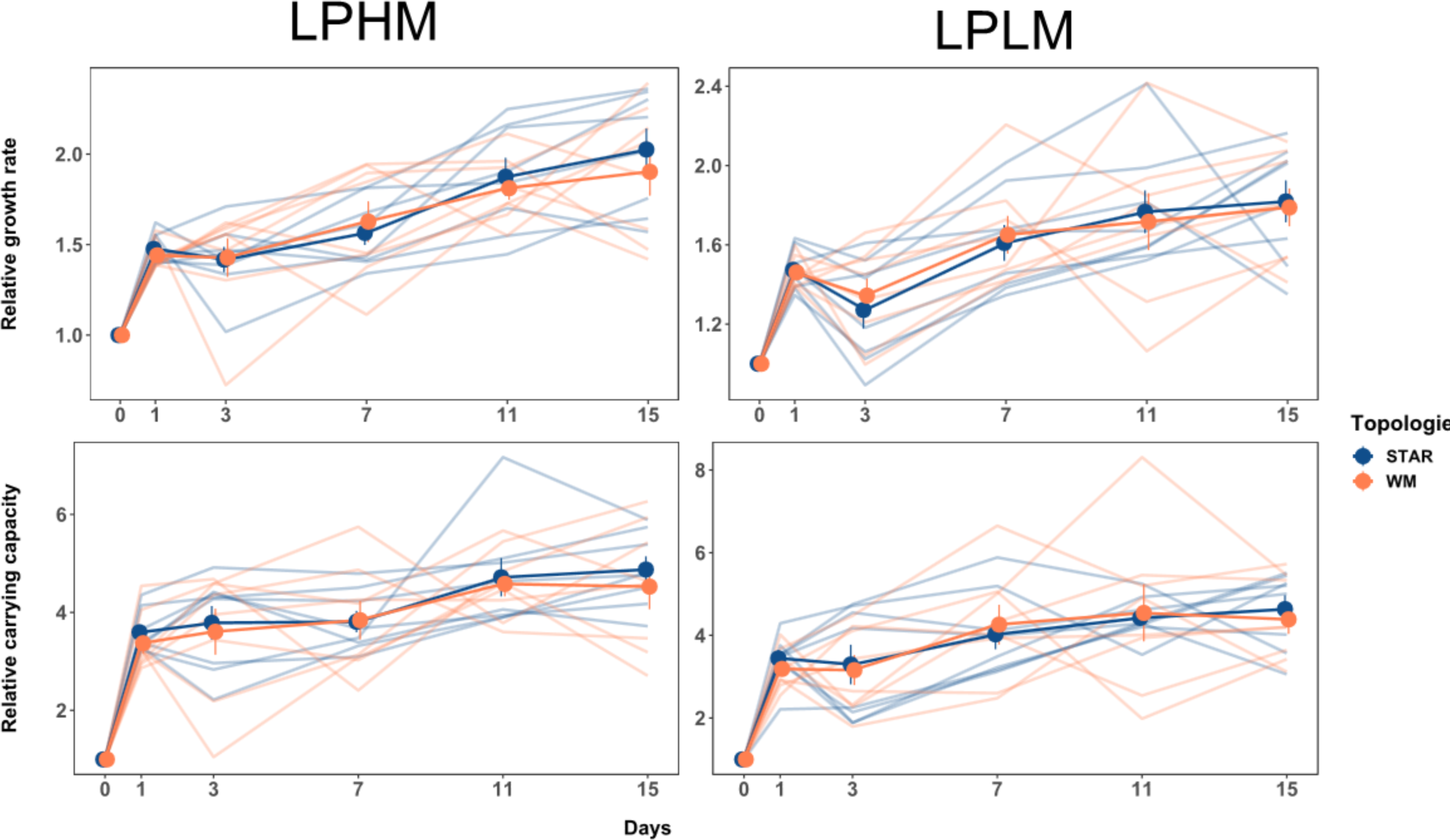
Dynamics of adaptation in large metapopulations. Fitness trajectories in large metapopulations connected by high and low migration rates (LPHM and LPLM). Increase in relative growth rate (upper panel) and relative carrying capacity (lower panel) in metapopulations propagated by either the star or the well-mixed topology with the LPHM (left panel) or LPLM (right panel) regime over the experimental time-period. Each point is the mean of 8 replicate metapopulations for a particular day and network topology, error bars show 1 standard error of the mean (SE). Raw data from each replicate metapopulation shown as faded lines. For large metapopulations (LPHM and LPLM), approximately 6.67 generations of growth happened per transfer.

Accelerated adaptation in low population size stars could result from the more rapid spread of one or a few large effect beneficial mutations or many more small effect mutations whose combined fitness is large, although given the short time frame of our experiment (∼100 generations) the latter explanation seems unlikely. To decide between these alternatives, we collected 48 clones from each of the evolved metapopulations and assayed stationary phase density at 24 hours, *K*, as a proxy for fitness in the selection medium (Figure 4). Large effect clones are preferentially fixed in the well-mixed population compared to the star when population sizes are large (Fig. 4 panel A-B, permutation KS test, p<10^-3^ and p<10^-3^, for high and low migration rates, respectively for 10000 permutations), consistent with strong clonal interference biasing fixation towards large effect mutants. In small populations, by contrast, star metapopulations harbored more large effect clones compared to the well-mixed metapopulations across both high and low migration rates (Fig. 4 panel C-D, permutation KS test, p<10^-4^ and p<0.05, for high and low migration rates, respectively for 10000 permutations). This result is hard to explain by clonal interference alone, which should be higher in well-mixed populations where there are more pathways for dispersal among subpopulations. Rather, we suspect that acceleration was caused by beneficial mutations being more likely to escape drift loss in stars than in well-mixed populations, consistent with the mechanism previously observed for a single, known beneficial mutation(*26*).

**Figure 3:**
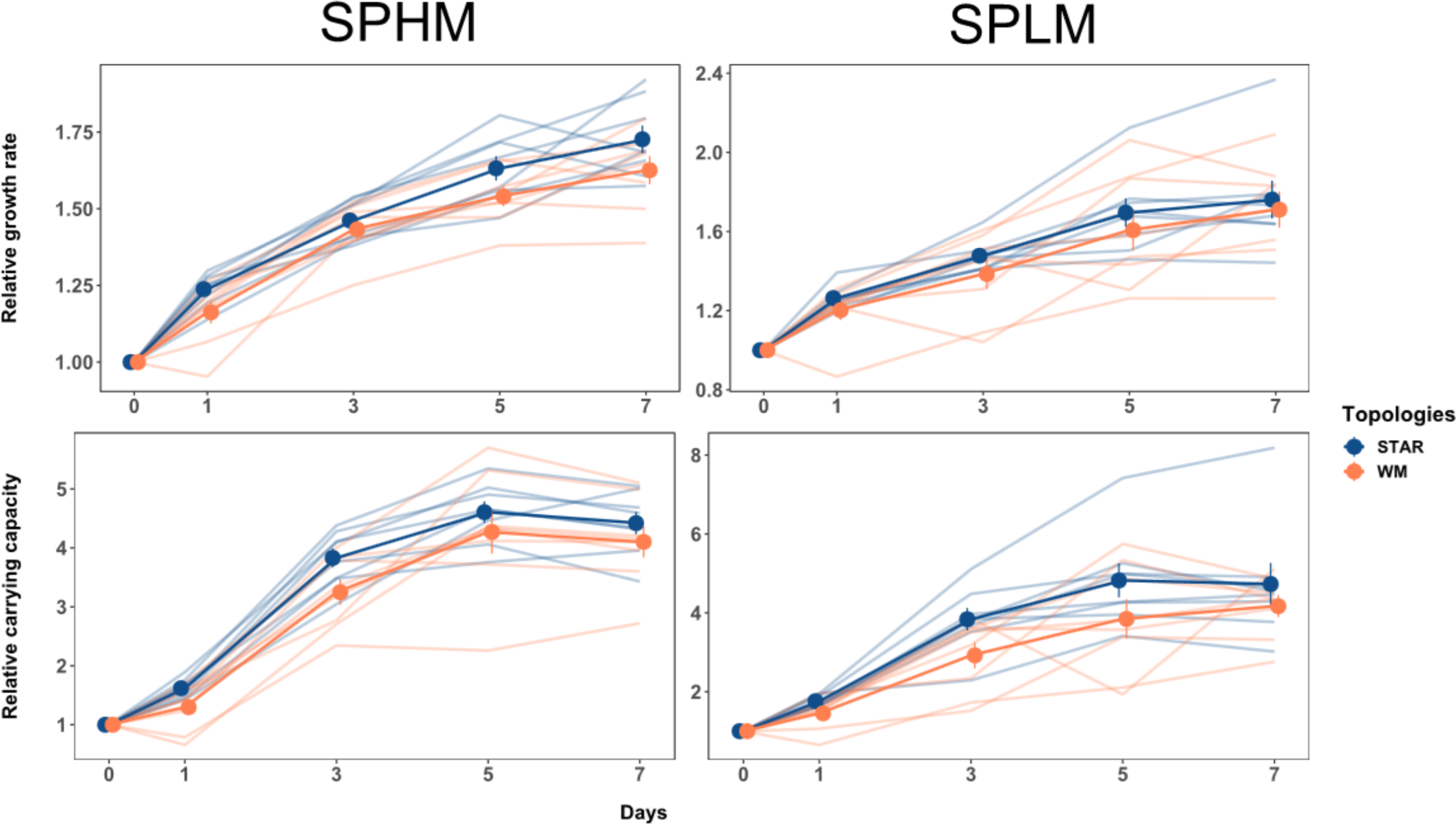
Dynamics of adaptation in small metapopulations. Fitness trajectories in small metapopulations (SPHM and SPLM) connected by high and low migration rates. Increase in relative growth rate (upper panel) and relative carrying capacity (lower panel) in metapopulations propagated by either the star or the well-mixed topology with the SPHM (left vertical panel) or SPLM (right vertical panel) regime over the experimental time-period. Each point is the mean of 8 replicate metapopulations for a particular day and network topology, error bars show 1 standard error of the mean (SE). Raw data from each replicate metapopulation shown as faded lines. Small metapopulations (SPHM and SPLM) experienced approximately 13.28 generations of growth per transfer.

**Figure 4:**
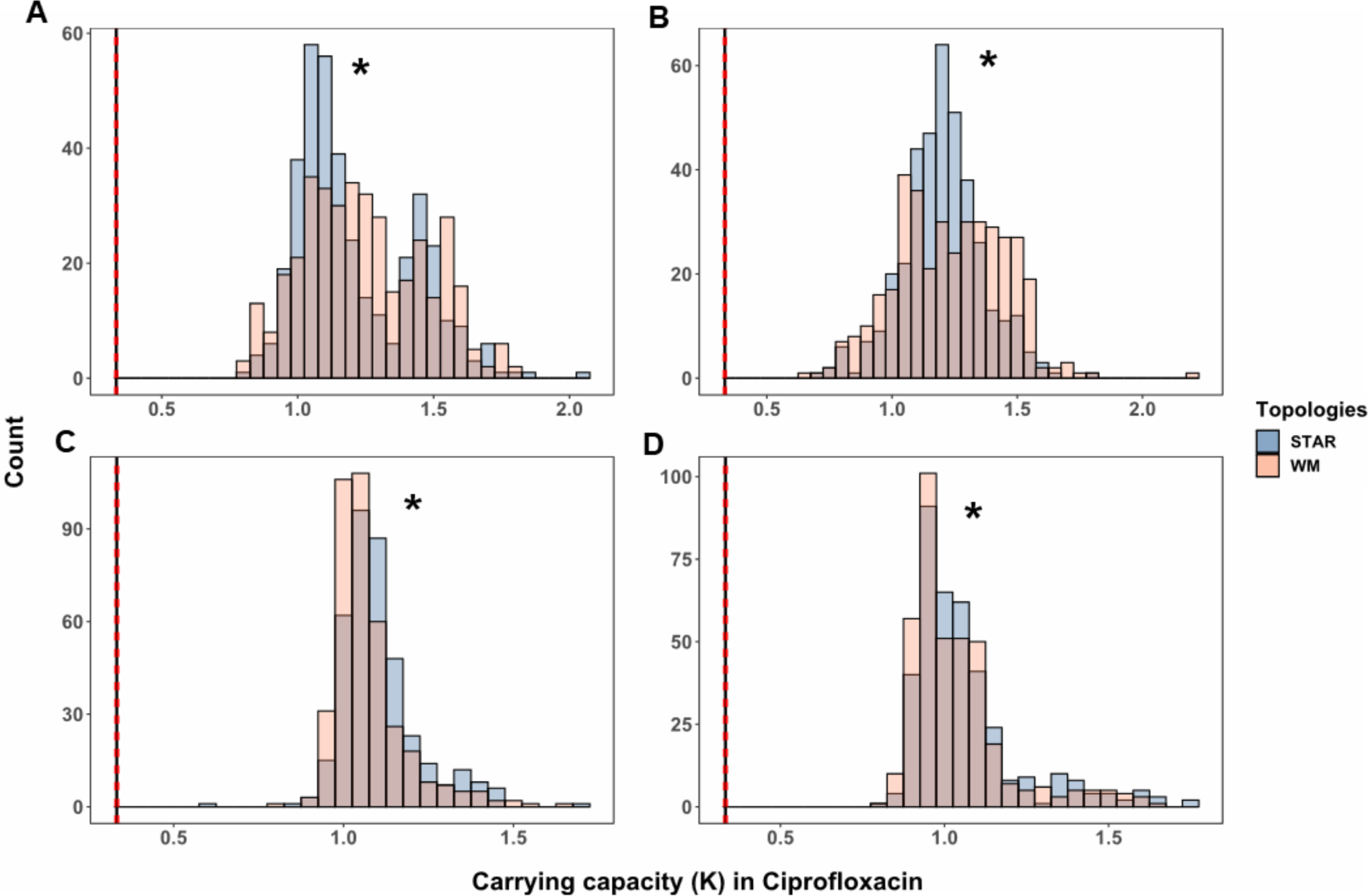
Distribution of fitness effects among isolates in the selection medium. Absolute fitness of evolved end-point isolates in the selection medium (LB supplemented with subinhibitory concentration of ciprofloxacin). Fitness effect distributions are shown as histograms of carrying capacity (*K*) for all mutants isolated from the large (LPHM and LPLM) metapopulations (A and B, respectively) or the small (SPHM and SPLM) metapopulations (C and D, respectively). Blue and red bars denote isolates collected from metapopulations propagated either by the star or the well-mixed topologies, respectively. The asterisk (*) indicates significance at P<0.05 for permutation K-S test (10000 permutations). The vertical line on the left-hand side of each histogram is the mean *K* of the ancestor in the selection medium and the red dotted vertical lines are one standard error (SE) of the mean.

Importantly, the reduction in the probability of drift loss only operates under low mutation supply rates. We checked that our experimental manipulations were effective at manipulating mutation supply rates by examining the number and spectrum of mutations segregating at the end of our experiment by sequencing 24 evolved metapopulations (6 for each population size x topology combination). We achieved ∼3000 fold sequencing depth and ∼98.5% genome wide coverage for each metapopulation which, after comparing the data with ancestral Pa14 genomes (Pa14 and Pa14-LacZ), allowed us to detect all mutations segregating above ∼ 5% frequency in each metapopulation. Figure 5 shows how many mutations on average segregate in each treatment. Large populations tend to harbour more mutations on average than small populations (2-way ANOVA, main effect population size, p=0.09), and well-mixed metapopulations support more mutations than well-mixed populations at both population sizes (2-way ANOVA, main effect of network treatment, p = 0.11). While these effects are marginally significant, they reassure us that our experimental manipulations effectively modulated the level of clonal interference in the expected directions.

**Figure 5:**
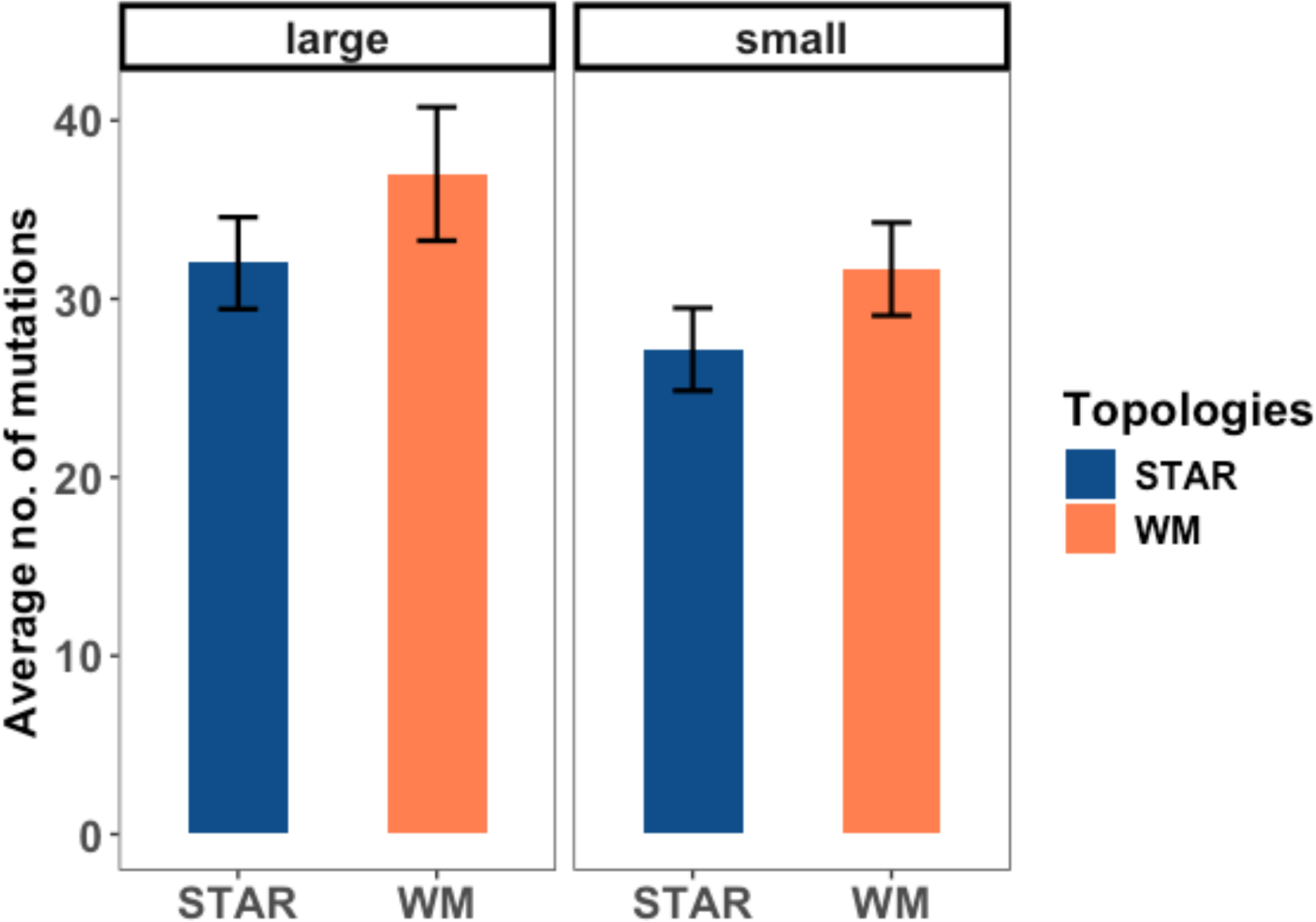
Number of mutational changes in each population size/network topology treatment combinations. Shown are the mean number of mutational changes for each network topology treatment (star and well-mixed) under each population size (large and small). Error bars represent one standard error of the mean.

If the cause of acceleration is the preferential substitution of large effect mutations due to their reduced probability of drift loss, then we should expect to see higher levels of repeated, or parallel, evolution in small *N* stars than in any other treatment of our experiment. Parallel evolution is often taken to be an indicator of strong selection acting in a particular locus, as it is unlikely that the same genes or mutations would fix repeatedly by chance alone(*28*). Equally, it could be due to preferential substitution caused by the reduced probability of drift loss, and so higher fixation probability, among beneficial mutations, in particular the rarer class of larger effect mutations available to selection(*28–32*). We tested this prediction by using our sequencing results to calculate the rate of repeated, or parallel, evolution for each mutated gene in our data set. We find that parallel evolution was higher in the small *N* star metapopulations compared with all others, an observation confirmed by the significant interaction between population size and network topology for three metrics of parallelism (p<0.05; see methods). Closer inspection reveals the two topologies do not differ in the extent of parallelism in large metapopulations but they do differ for small metapopulations, stars showing significantly higher parallelism than their well-mixed counterparts (Table 1). As there is little reason to suspect the distribution of fitness effects among beneficial mutations differs systematically between star and well-mixed topologies for a given population size, the high levels of parallelism in the small *N* star must be associated with a higher fixation probability among the first mutations arising in the experiment. Given the over-representation of large effect beneficial mutants in small *N* star metapopulations, these results suggest accelerated adaptation is due to large effect beneficial mutations having a higher probability of fixation because they avoid drift loss when rare in stars compared to well-mixed populations.

**Table 1:** Population-level parallelism. Difference in population-level parallelism between the two topologies for each effective population size. Differences between the mean values for each metric (estimate) in the star and the well-mixed topologies are presented. A positive value denotes the calculated metric for the star topology is higher than the well-mixed topology and vice versa. Significance (P<0.05) is determined by a 2-way ANOVA for dispersion and a 2-way ANOVA followed by a permutation test (10,000 permutations) for Jaccard distance and C-score (materials and methods).

|  | Estimate = (Star - Well-mixed) |  |  |  |  |  |
| --- | --- | --- | --- | --- | --- | --- |
| Population size | Dispersion |  | Jaccard Distance |  | C-score |  |
|  | Estimate | Significance | Estimate | Significance | Estimate | Significance |
| Large | 0.0167 | 0.7153 | 0.0165 | 0.6569 | 0.0219 | 0.5356 |
| Small | -0.1265 | **0.0108** | -0.1514 | **0.0009** | 2.0156 | **0.0029** |

As expected from the strong selection pressure exerted by ciprofloxacin, we found many known resistance mutations circulating in moderate to high frequencies in these metapopulations (Figure 6). We identified frequent mutations (small indels and nonsynonymous SNPs) in the negative regulators and components of efflux pumps such as mexA and mexR (mexAB-oprM), mexS (MexEF-OprN) and nfxB (MexCD-OprJ) that are constitutively expressed when it is necessary to decrease intracellular concentration of ciprofloxacin(*33, 34*). We also uncovered mutations that prevent ciprofloxacin from binding to the DNA-modifying subunits of DNA gyrases (gyrA, gyrB) (*35–37*). Not unexpectedly, our evolved metapopulations also harbor mutations in genes that do not confer resistance to ciprofloxacin but are global regulators of quorum sensing (lasR), c-di-GMP signaling and biofilm formation (wspA/F/R, morA), virulence and twitching motility (pilB/C/D/F)(*38–51*). We and others commonly observe these same mutations in rich laboratory growth media, where their fitness advantage is thought to be associated with reducing the costs of metabolism(*52, 53*).

**Figure 6:**
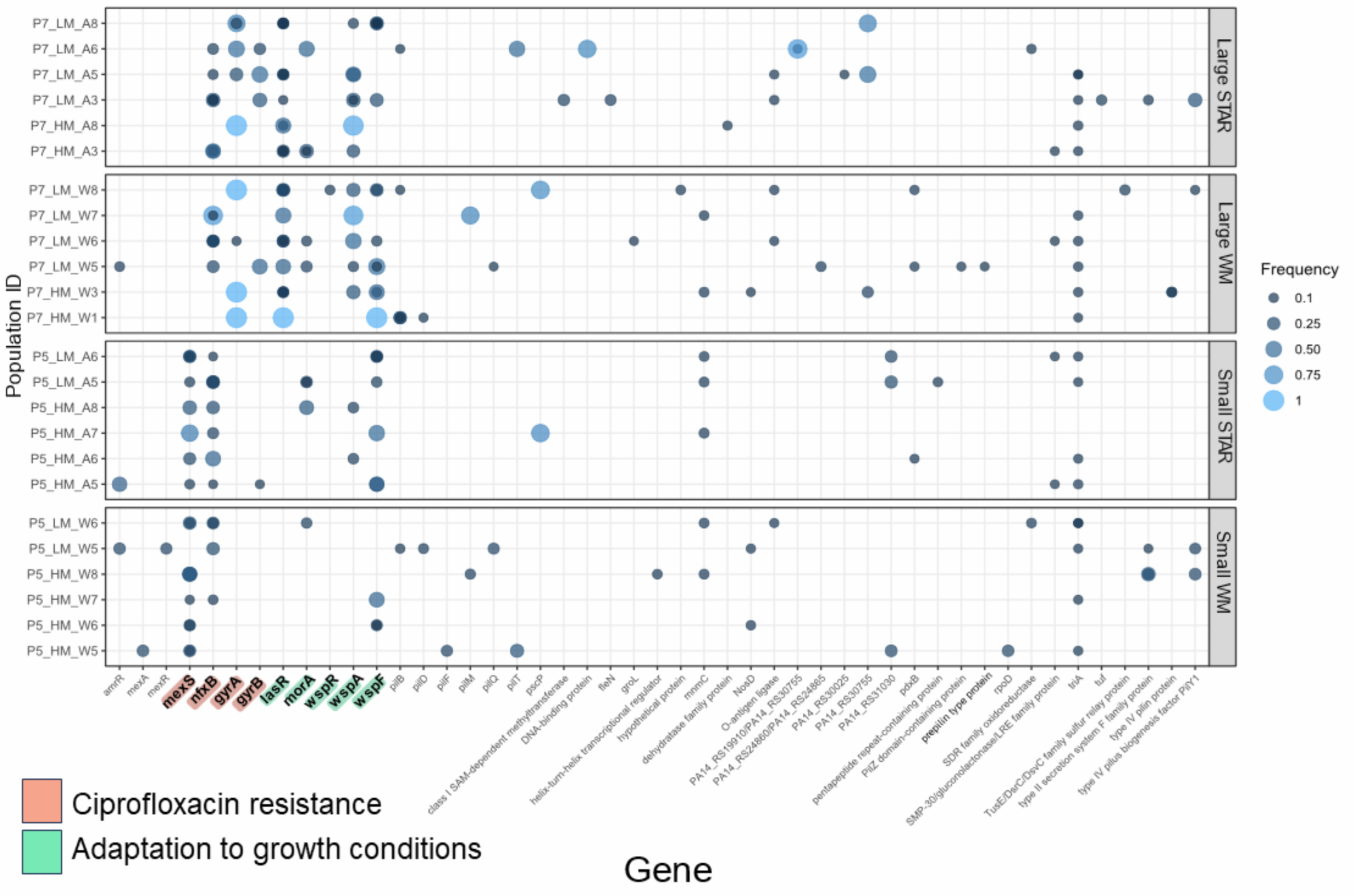
Genetic changes detected in the metapopulations. (not showing intergenic mutations; full data file is available in supplementary figure 1) after ∼ 100 generations of evolution in the selection media (LB supplemented with subinhibitory concentration of ciprofloxacin). From the top, the first and the second panel show large star and large well-mixed metapopulations, respectively. Similarly, the third and the fourth panel show small star and small well-mixed metapopulations, respectively. Mutation in genes highlighted with either red or green background denote recurring canonical ciprofloxacin resistance genes and genes that are relevant for adaptation of *P. aeruginosa* to laboratory conditions, respectively. Size of the circles depict observed frequencies of mutants in the metapopulations. More than one circle for a gene represents the presence of distinct genetic variants (alleles) in the same metapopulation.

A last striking result from our genomic analyses is that metapopulations evolved under high and low mutation supply diverge in the repertoire of resistance mutations they accumulate over the evolutionary period. Notable examples include the enrichment of *mexS* and *gyrA* mutations exclusively in small and large metapopulations, respectively (binomial test, p<0.005 and p<0.05, respectively). We see the same pattern for putative non-resistance mutations as well: *lasR* and *wspA* being specifically mutated in the large (binomial test, p<0.005 and p<0.05, for lasR and wspA, respectively) but not in the small metapopulations. Although we do not uncover any mutations that are network specific in the large metapopulations there is a marginally significant enrichment of mutations in *wspA* and *wspF* specific to the small star metapopulations (*wspA* and *wspF* mutations in 6 star compared to *wspF* mutations in 2 well-mixed metapopulations). Additionally, it is notable that mutations in *nfxB* are present in all (6 out of 6) small star but only 50% (3 out of 6) in the small well-mixed metapopulations sequenced, although the difference is not statistically significant (binomial test, p=0.08). Taken together, this data is consistent with the idea that mutation supply rate can bias the spectrum of mutations contributing to adaptation. Large populations with high mutation supply rates have access to a larger spectrum of mutations and clonal interference during substitution can lead to preferential fixation of those with the largest fitness effects. Interestingly, however, there is little evidence that network topology itself biases the spectrum of mutations available to selection; rather, it is the preferential enrichment of large effect mutations that avoid drift loss in star metapopulations that governs the dynamics of molecular variation in these situations.

## Discussion

Our work provides direct, experimental evidence that the topology of connections in a metapopulation markedly alters the pace of adaptive evolution and the identity of the mutations responsible. Specifically, we have shown that small *N* star metapopulations, those characterized by satellite populations connected through a central hub, can accelerate adaptation over and above that observed in a well-mixed setting. Acceleration is associated with an enrichment of large effect mutants and results in higher levels of gene-level parallelism compared to well-mixed populations of the same size. These results are consistent with an underlying mechanism involving the concentration of beneficial mutations in the hub by dispersal, reducing their probability of drift loss and facilitating their rapid spread to other leaves in the metapopulation.

That we observed acceleration in the small *N* stars is surprising because it does not fit with the two main models of reproduction used to track allele frequencies in metapopulations. The approach most common in population genetics, the Fisher-Wright model, assumes reproduction involves bulk replacement of parents by offspring in non-overlapping generations. Metapopulation topology has little or no effect on fixation probabilities in this model, and the pace of adaptation is almost always fastest in well-mixed populations(*9, 10*). An alternative approach, evolutionary graph theory, is based on the Moran model of individual reproduction and death, which leads to overlapping generations. Moran-based models show that stars can increase the fixation probability of beneficial mutants relative to a well-mixed population, although at the cost of longer fixation times(*5, 7, 13*). Our results show that both models fail to capture accurately the dynamics of adaptation on stars: in contrast to Fisher-Wright models stars can adapt faster than well-mixed populations whereas times to fixation appear to be much smaller than anticipated for the Moran process on graphs. Clearly, both models need revision.

How common is accelerated adaptation on star-like structures in more natural systems? This remains difficult to answer. Many microbial communities occupy environments such as soil or hosts that are spatially structured(*54–61*), and some studies have shown that spatial structure following population bottlenecks imposed during host dissociation or biofilm dispersal can have profoundly different outcomes depending on the spatial structure of the population(*56, 62, 63*). The fact that we only observed acceleration at the smallest population sizes in our experiment (*N_e_* ∼ 10^5^ individuals) suggests there could be an upper threshold of population size above which acceleration is unlikely to be observed. Nevertheless, the metapopulation structure of many large organisms may resemble that of a star, for example when range edge populations are sparse compared to the more abundant center or due to habitat fragmentation(*2, 3*). If so, star topologies could be an explanation for the rapid spread of invasive species. Moreover, many scale-free networks, which often characterize contact networks in epidemiology, contain sub-structures resembling stars and are known to be weak amplifiers of selection(*1, 13*). It is not inconceivable that metapopulation structures such as the star play a more important role in adaptive evolution than has been recognized up to now.

## Methods

### Microbial strains and conditions

For all experiments, clonal populations of *Pseudomonas aeruginosa* strain 14 (PA14) and PA14:lacZ, isogenic to PA14 except with an insertion in the lacZ gene respectively, were used. Colonies possessing the lacZ insertion appear blue when cultured on agar plates supplemented with 40 mg/L of 5-bromo-4-chloro-3-indolyl-beta-D-galactopyranoside (X-Gal), and are visually distinct from the PA14 white colouration. The neutrality of the lacZ marker was confirmed in our experimental environments by measuring the fitness of the marked strain relative to the unmarked strain. Populations were cultured in 24-well plates with 1.5 mL of media in each well, in an orbital shaker (150 RPM) at 37°C. The culture media consisted of Luria Bertani broth (LB: bacto-tryptone 10 g/L, yeast extract 5 g/L, NaCl 10 g/L) supplemented with 40 ng/mL of the fluoroquinolone antibiotic, ciprofloxacin. This particular subinhibitory concentration of ciprofloxacin was chosen to exert a moderate level of selection that would slow down the dynamics of fixation of resistant mutations compared to a lethal dose above the minimum inhibitory concentration (MIC). This approach allows us to discern the effect of network topology on the process of adaptive substitution with weakened interference of strong selection (supplementary figure 7, *26*). Lower ciprofloxacin concentrations did not enrich resistant mutations to high levels in the populations within the experimental time period tested (supplementary figure 8, *26*). All strains and evolving populations were cryopreserved at −80°C in 20% (v/v) glycerol.

### Evolution experiment

A single metapopulation consisted of four subpopulations, one subpopulation being located on each of four different 24-well plates. Plate 2 was always assigned as the hub, and plates 1, 3, and 4 were treated as the leaves. Half of the metapopulations (every odd numbered replicate) in this experiment was inoculated with clonal PA14 and the other half (every even numbered replicate) with clonal PA14:LacZ. This was necessary in order to track any cross-contamination between metapopulations inhabiting adjacent wells in the 24 well plate.

In a metapopulation, mutation supply rate can be manipulated at two different levels, at the level of subpopulations -which can be manipulated by changing the effective population size (*N_e_*) of the constitutive subpopulations and at the level of the whole metapopulation - by modifying the rate of migration (*m*) between the subpopulations. In our experiment, during daily serial transfer of the metapopulations we manipulated the mutation supply rate by creating a total of four combinations of effective population size (large and small) and migration rate (high and low, relative to the effective population size). For each of these mutation supply rates, replicate metapopulations were propagated by either the star or the well-mixed network topologies. This full-factorial design allowed us to track a total of 64 replicate metapopulations (8 replicate metapopulations X 2 topologies X 4 mutation supply rates). MIgration in the star was biased towards the hub (3X In>OUT) as previous work showed this pattern of asymmetric migration resulted in faster spread of a single beneficial mutation than symmetric bidirectional dispersal (*26*).

Experiments were initiated by inoculating each subpopulation in a metapopulation with ∼10^7^ (large) or ∼10^5^ (small) colony forming units (CFU) per ml of either PA14 or PA14:lacZ descended from a single colony picked from an agar plate and grown overnight in liquid LB at 37°C with vigorous shaking (150 RPM). Metapopulations were transferred daily following dispersal among subpopulations (see below) by diluting overnight cultures either 1:10^2^ or 1:10^4^ into fresh medium supplemented with ciprofloxacin. This transfer regime corresponds to ∼6.67 (large) or ∼13.28 (small) daily generations of growth (N_t_ = N_0_ X 2*^g^*, N_t_/N_0_ = 10^2^ or 10^4^, *g* = number of generations). In our experiments, each subpopulation of a large metapopulation had a ∼50 times higher effective population size (N_e_ = N_0_ X *g*) than that of a small metapopulation.

We constructed distinct network topologies by mixing subpopulations prior to serial transfer following the schematic shown in our previous work (supplementary figure 4, *26*). Briefly, well-mixed networks were created by combining equal volume aliquots from all subpopulations into a common dispersal pool, diluting this mixture to the appropriate density to achieve the desired migration rate, and then mixing the dispersal pool with aliquots from each subpopulation (so-called ‘self-inoculation’) before transfer. Star networks, which involve bidirectional dispersal between the hub and leaves, were constructed in a similar way to the well-mixed situation only now the dispersal pool consisted of aliquots from just the leaves and aliquots from the hub (plate 2) were mixed with ‘self-inoculation’ samples from each leaf prior to serial transfer. For LPHM, LPLM, SPHM and SPLM ∼10^5^, ∼10^3^, ∼10^3^, ∼10 CFU/ml migrants were used in addition to the self-inoculations, respectively. Further details on how each network topology and migration rate were achieved are the same as ref (*26*).

This daily transfer protocol was continued for ∼100 generations, corresponding to 15 transfers (“days”) for the large metapopulations and 7 transfers (“days”) for the small metapopulations.

### Phenotypic analyses

#### Measurement of fitness of the metapopulations

We measured the fitness of each evolved metapopulation every ∼25 generations throughout the evolution experiment. We used growth rate (*r*) and carrying capacity (*K*) in the selective medium as proxies for fitness. During the evolution experiment, a mixture of the whole metapopulation was archived daily. We revived these metapopulation mixes saved on Day 1, 3, 7, 11 and 15 for the large metapopulations, and Day 1, 3, 5 and 7 for the small metapopulations in the selection medium by overnight growth in 1.5 mL of media in each well, in an orbital shaker (150 RPM) at 37°C along with the PA14 and PA14:LacZ ancestors. On the next day, the overnight cultures were diluted 1:1000 in fresh 200μl selection medium in a 96 well plate and a growth curve experiment was started in a BioTek PowerWave spectrophotometer (BioTek Instruments Inc., Winooski, VT) incubated at 37°C with linear shaking for 20 seconds every 5 minutes. The optical density (OD) of each sample was measured at 600 nm after every 20 minutes for 24 hours until the sample reached stationary phase. Carrying capacity was measured as the highest optical density reached during the 24 hour growth and growth rate was estimated as the maximum slope of the growth curve with a procedure described in another study (*64*). Each measurement was carried out with five biological replicates. We only revived cryo-stocks belonging to the same day together and the measured carrying capacity and growth rate was normalized by the appropriate ancestors’ growth on the same day of experiment, hence minimizing any measurement bias. Border wells of the 96 well plate were uninoculated to minimize sample loss due to evaporation.

#### Measurement of fitness of the evolved isolates collected from the metapopulations

We collected 12 single isolates from each subpopulation from all the evolved metapopulations (12 isolates X 4 subpopulations X 64 metapopulations = 3072 single isolates in total). The evolved subpopulations from the last time-point in the experiment were diluted 1:10^6^ and plated on LB-agar plates for overnight growth and inoculated at 37°C. Twelve single isolates from pre-marked positions were collected to avoid bias and were grown overnight in liquid LB medium and were cryo-preserved. The cryo-stocks for each isolate were streaked on a LB-Agar plate and a single colony was grown in the selective growth medium in 24 well plates overnight at 37°C with vigorous shaking (150 RPM). The optical density (OD) was measured at 600 nm in a BioTek PowerWave spectrophotometer after thoroughly mixing the overnight culture. For each isolate, carrying capacity was measured only once given the large number of total isolates for measurement. However, PA14 and PA14-LacZ ancestors grown 64 times each for CIP and NO-CIP environments suggests there was negligible day to day variation in carrying capacity measurements even for biological replicates (vertical red dashed lines in Figure 4 and 5).

### Genomic analyses

#### Whole-Genome Sequencing

We randomly selected 6 replicate end-point metapopulations from each treatment combination (2 population sizes (large or small) X 2 topologies (star or well-mixed) X 6 replicates = a total of 24 metapopulations). Among the selected metapopulations, for each topology two were from LPHM, 4 from LPLM, 4 from SPHM and finally 2 were from SPLM (Figure 1). The rationale behind this selection was two-fold: (a) LPLM and SPHM were seen to have the biggest differences between the topologies in the phenotypic analyses; and (b) the different migration rates did not influence the fitness trajectories of the metapopulations.

Both ancestral strains of *P. aeruginosa*, PA14 and PA14-LacZ, were also sequenced to facilitate genome assembly and to identify genetic variants arising during the evolution experiment. The four constituent subpopulations of each selected metapopulation were revived overnight from frozen stock and each metapopulation mix reconstructed by mixing the revived subpopulations in equal volumes. Genomic DNA was then extracted from each metapopulation mix for whole-genome sequencing using the QIAGEN DNeasy UltraClean 96 Microbial kit, following the manufacturer’s recommended protocol. Library preparation and sequencing were performed by Genome Quebec at McGill University on the Illumina NovaSeq 6000 platform, using paired-end sequencing of 2 × 150 base-pair reads.

#### Processing of genomic data and variant calling

Whole-genome sequencing of 24 metapopulations yielded a total of ∼750 Gb of raw data, with a median depth of 3622.3-fold and an average of 98.5% genome coverage. Sequencing reads were first quality checked by generating FastQC reports using FastQC version 0.11.9 and quality trimmed using Trimmomatic version 0.39 (Bolger et al. 2014), with the command SLIDINGWINDOW:5:20 MINLEN:20 LEADING:5 TRAILING:5 CROP:140 HEADCROP:10. Variants were called using Breseq version 0.36.1 (Deatherage and Barrick 2014) - a tool specifically designed for detecting mutations in microbial genomes, with default parameters (detection limit of 5%) and -p flag for detecting polymorphisms in the sequenced metapopulations. Reads were aligned to the *P. aeruginosa* reference genome UCBPP-PA14 - assembly GCF_000014625.1(*65*). We subsequently discarded variants common across both ancestral strains (PA14 and PA14-LacZ) and all evolved populations using the gdtools module of breseq to identify only the mutations that arose over the course of the selection experiment. We found no evidence of cross-contamination in our sequenced metapopulations (each even numbered replicate had the LacZ insertion which was absent in the odd numbered replicates). Also, none of the metapopulations harbored any mutations in the genes that have been linked to increased mutation rates(*66–68*) in *P. aeruginosa*, so we did not have to discard any sequenced genomes from further downstream analyses. All genomic analyses were performed on Compute Canada high performance computing platform using a custom bash script for bioinformatic workflow. Breseq output files with information on genetic changes were processed further for performing statistical tests and generating plots in R statistical computing software.

### Statistical analyses

All statistical analyses were performed in R (version 4.2.2)(*69*).

We modeled the rate of change in fitness (both relative growth rate and carrying capacity) through time separately for each population size treatment using a linear mixed model (lmer) with time(*70*), migration rate and network topology as fixed factors, and a random intercept for individual replicate population to account for resampling across time (repeated measures). Time was considered to be a second order polynomial regressor in our model since the trajectories were not linear. The distribution of fitness effects among the beneficial mutants isolated from the end-point metapopulations were compared between the network topologies using a Kolmogorov-Smirnov (K-S) test paired with a resampling procedure (10000 permutations).

To quantify gene-level parallelism in our experiment we focused on genes reaching or exceeding a frequency of 8% and excluded synonymous mutations under the assumption they are selectively neutral (supplementary fig. 2, since ∼97% synonymous mutations reached the frequency of ∼8% or less in our experiment). Since there is no broadly accepted metric for quantifying gene-level parallelism, we use three distinct measures: variance in dispersion of Euclidean distances between populations, Jaccard distance (Jdist = 1 - Jaccard Index, which describes the likelihood that the same gene is mutated in two independent populations) and observed repeatability relative to expectation under randomness using the hypergeometric distribution (C-score) (*sensu 53*). Briefly, a low value for dispersion, a low Jaccard distance and a high C-score indicates a high degree of parallelism and vice versa.

For dispersion, we calculated the mean distance between a population and the corresponding population or network topology treatment centroid, following a PCoA on a Euclidean distance matrix using the vegdist function from the vegan package in R(*71*). The distance is measured as the population and network topology level genetic variance with larger mean dispersion signifying more divergent genetic changes and therefore less parallelism.

For the Jaccard distance measure, we calculated the Jaccard measure from the vegdist function from the vegan package in R as the dissimilarity between all pairs of populations within a treatment (effective population size or network topology)(*53*). Jaccard distance is the complement to Jaccard index and describes the likelihood that the same gene is mutated in two independent populations (here metapopulations). Jaccard distance can range 0 to 1: zero being two samples are exactly alike and one being two samples are completely different. We reported the difference between the network topology means (estimated difference = star - well-mixed, averaged over samples) for each effective population size treatment. For the J-distance, a positive difference denotes higher parallelism for well-mixed and negative difference means higher parallelism for the star topology.

Our last measure of repeatability is the C-score, which uses the hypergeometric distribution to calculate the deviation between the observed amount of parallelism and the expectation under random gene use(*72*). The magnitude of the C-score represents the magnitude of the deviation, larger C-scores signifying higher repeatability and therefore more parallelism. As for the Jaccard distance we report the differences between the C-score means for each treatment combination, a positive difference denotes higher parallelism for a star and negative difference means higher parallelism for the well-mixed topology.

To determine the significance of each metric, we first performed an 2-way ANOVA with the metric (Euclidean distance, Jaccard distance or C-score) as the response variable and effective population size and network topology and their interactions as explanatory variables. We followed up any significant interaction between effective population size and network topology by measuring the difference in gene-level parallelism between the two topologies for each effective population size (significance threshold set at p<0.05 for all three metrics) using the emmeans package from R(*73*). Furthermore, to determine the significance of both Jaccard and C-score metrics, we performed a permutation test by randomizing population size and network topology treatment labels (number of permutations = 10,000) and calculating a null distribution of F-values. We complemented this analysis by calculating the probability (out of the total number of permutations) of the observed difference between the star and the well-mixed topologies being higher (C-score) or lower (J-distance) than the randomized estimated difference between the topologies, for each effective population size.

Parallelism at the gene level was defined as the proportion of populations with mutations in that gene, both globally for effective population size and network topology and within all of the four treatment combinations(*53*). To test for significance, we calculated the probability of our observed results against the null hypothesis that gene use was random, using the binomial distribution with the number of metapopulations as the number of trials, number of times a gene was mutated as the number of successes, and proportion of total metapopulations across all treatments with a mutation in that gene as the probability of success. From this, if the probability of an observation was <0.05, we considered that gene to be either effective population size or network topology treatment specific.

## Acknowledgement

This work was supported by a Natural Sciences and Engineering Research Council (NSERC) Discovery Grant to RK.

## Author Contribution

R.K. conceptualized the project and provided guidance on lab techniques and analyses. P.P.C. performed the experiments and analyzed the data. P.P.C and R.K. wrote the manuscript.

## Materials & correspondence

All correspondence and requests for materials should be directed to Rees Kassen.

## Data availability

All raw data generated, genome sequences and R scripts used for data analyses will be made available online upon publication.

