## Supplementary figures and images for "Rapid adaptation and increased genetic parallelism in experimental metapopulations of *Pseudomonas aeruginosa*"

### Supplementary figure 1

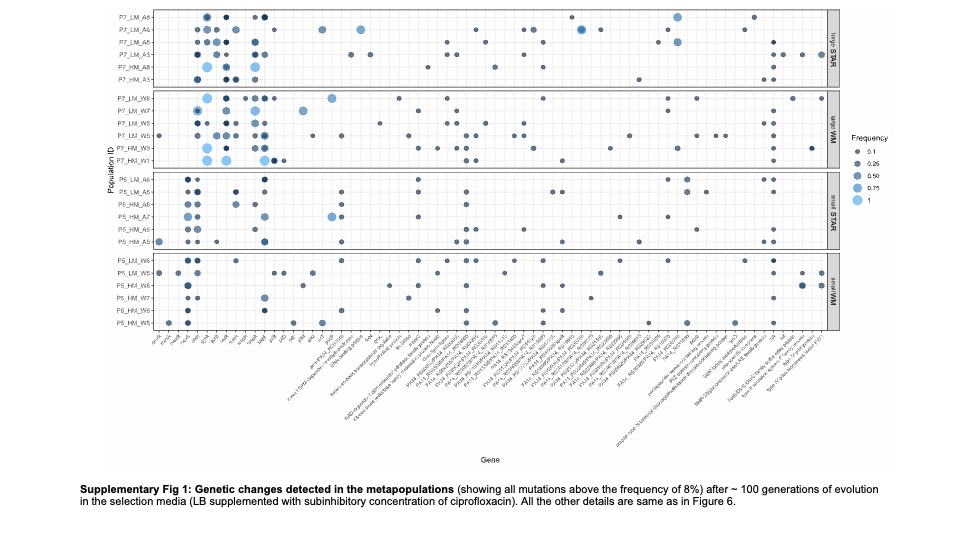

### Supplementary figure 2

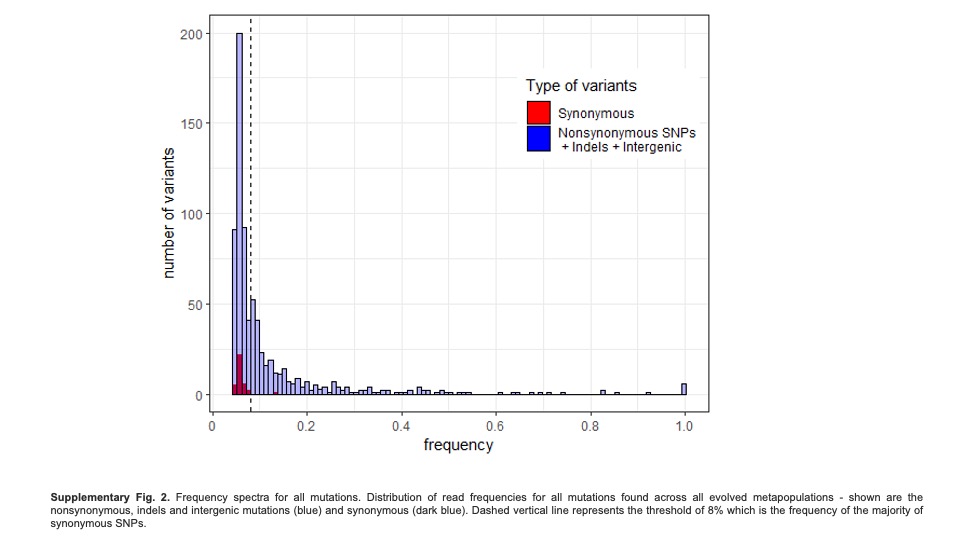
